# Neural Correlates of Causal Confounding

**DOI:** 10.1101/395707

**Authors:** Mimi Liljeholm

## Abstract

As scientists, we are keenly aware that if putative causes perfectly co-vary, the independent influence of neither can be discerned – a “no confounding” constraint on inference, fundamental to philosophical and statistical perspectives on causation. Intriguingly, a substantial behavioral literature suggests that naïve human reasoners, adults and children, are tacitly sensitive to causal confounding. Here, a combination of fMRI and cognitive computational modeling was used to investigate neural substrates mediating such sensitivity. While being scanned, participants observed and judged the influences of various putative causes with confounded or non-confounded, deterministic or stochastic, influences. During judgments requiring generalization of causal knowledge from a feedback-based learning context to a transfer probe, activity in the dorsomedial prefrontal cortex (DMPFC) was better accounted for by a Bayesian causal model, sensitive to both confounding and stochasticity, than a purely error-driven algorithm, sensitive only to stochasticity. Implications for the detection and estimation of distinct forms of uncertainty, and for a neural mediation of domain general constraints on causal induction, are discussed.

## Introduction

Consider two scenarios involving a target cause, *C*, an alternative cause, *A*, and the presence (+) or absence (-) of some effect. In the first scenario, the effect occurs when *C* and *A* are presented in combination, but not when *A* is presented alone [*A*-, *AC*+]. In the second scenario, the *A*-trials are removed, so that both *C* and *A* occur only in combination with each other [*AC*+]. How would your judgment about the influence of *C* differ across these two scenarios? Philosophers, scientists and statisticians alike recognize that perfect co-variation of *C* and *A* in the second scenario occludes the independent influences of each putative cause, rendering any judgment about their respective effects fraught with uncertainty. Importantly, naïve human reasoners also appear to, tacitly, apply this domain-general constraint on causal induction, as evidenced by behavioral data showing sensitivity to confounding in a wide range of causal and predictive judgments, by children as well as adults (Spellman, 1996; Kushnir and Gopnik, 2005; Meder et al., 2006; Schulz and Bonawitz, 2007; Schulz et al., 2007; Liljeholm, 2015). Very little is known, however, about the neural substrates mediating sensitivity to, and uncertainty associated with, causal confounding. The current study aims to identify such neural computations.

A second source of uncertainty in causal and predictive inference, that has been thoroughly investigated both behaviorally and neutrally, is the stochasticity, or variance, of the outcome variable, which is greatest when the probability distribution over possible outcome states is uniform. As with causal confounding, behavioral sensitivity to outcome stochasticity is well established (e.g., Holt & Laury, 2002). Moreover, several neuroimaging studies have implicated the dorsomedial prefrontal cortex (DMPFC), anterior insula, thalamus and dorsolateral prefrontal cortex (DLPFC) in the neural representation of outcome stochasticity (Volz et al., 2003; Huettel et al., 2005; Grinband et al., 2006; Abler et al., 2009). An important open question is whether, and which of, these neural regions also implement signals associated with causal confounding. In the current study, participants were scanned with functional MRI while observing and judging the influences of various putative causes with confounded or non-confounded, deterministic or stochastic, influences.

Note that, if a particular set of confounded causes always occur in the same configuration, and the only goal is to predict the state of the outcome based on that particular configuration, then confounding does not pose a problem. Indeed, most individual, non-confounded, causes can presumably themselves be broken into a set of always co-occurring, and thus confounded, elements. The critical question, thus, is whether it is necessary to tease apart the influences of individual elements, such as when a confounded cause suddenly occurs on its own, or in a novel combination with some other cause. In the current study, participants were required both to make predictions about the outcome given a particular, recurring, configuration of confounded causes and, in other instances, to make explicit judgments about the individual influences of those same causes. Only in the latter case would uncertainty due to confounding be warranted.

Formally, two dominant approaches to predictive uncertainty can be discerned in the psychological literature. First, in associative learning theory, an error-driven representation of uncertainty about a cue’s predictive strength is captured by the “Pearce-Hall” (PH) algorithm, which relates the “associability” of a cue to a weighted average of the absolute prediction error on previous trials involving that cue. Since the frequency and size of prediction errors increases with unpredictability, this quantity is proportional to the stochasticity of the outcome. Conversely, in Bayesian Causal Models, uncertainty about the predictive strength of a cause is reflected in the entropy of the posterior distribution over its possible strengths, which depends on the variance of the effect variable, but also on the independent occurrence of alternative causes. While both the associative and Bayesian causal model predicts sensitivity to stochasticity, only the causal model accounts for uncertainty due to causal confounding. Here, a combination of neuroimaging and cognitive modeling was used to dissociate neural signals scaling with error-driven and causal uncertainty. In particular, judgments requiring generalization of causal knowledge from a feedback-based learning context to a transfer probe were expected to elicit strong neural responses to both stochasticity and confounding.

## Materials & Methods

### Participants

20 healthy normal volunteers (mean age = 20.9 ± 2.4, range: 18-27, 12 females) participated in the study. A power analysis performed on data from a pilot study on error-driven uncertainty, described in detail below, indicated that a sample size of 16 would yield a power of 0.9 at a Gaussian random field theory corrected threshold of 0.05 in regions of interest. One participant was excluded prior to any analyses due to excessive head movement (> 6mm), leaving a sample size of 19. The volunteers were pre-assessed to exclude those with a history of neurological or psychiatric illness. All subjects gave informed consent and the study was approved by the Institutional Review Board of the University of California, Irvine.

### Task & procedure

Participants were scanned with functional MRI while performing a causal induction task in which they assumed the role of a research scientist assessing the influence of various allergy medicines on headache – a potential side effect. At the beginning of the study, participants were instructed that each medicine could either produce headache or have no influence on headache (i.e., there were no preventive causes) and, further, that the influence of a given medicine might be stochastic, so that even if that medicine was indeed a cause of headache, it may still not produce headache every time it was administered. Three target medicines, C, D and S, mnemonically labeled here to indicate “confounded”, “deterministic” and “stochastic” influences respectively, occurred only in combination with some alternative medicine during feedback-based learning. Specifically, with +, - and ± respectively indicating a 1.0, 0 and 0.5 probability of the effect, and with lower case letters indicating non-target causes, there were 5 different types of medicine treatments: e-, s’±, De+, Se± and Cc’+. None of the allergy patients had headache in the absence of any medicine, which was explicitly stated in the initial instructions as well as apparent on several “no medicine” (n-) trials.

Note that medicines C and c’ both occurred *only* in combination with one another, so that target medicine C was perfectly confounded with its compound counterpart. Medicine D also occurred only in combination with another medicine, e, however, “elemental” medicine e also occurred by itself, allowing for an estimation of the independent causal influence of medicine D across the constant presence of e (i.e., across e- and De+ trials). Medicine S was identical to medicine D, except that the probability of headache given the Se compound was 0.5 rather than 1.0; a comparison of medicines S and D, therefore, identifies a difference between deterministic and stochastic causation. Finally, medicine s’ also produced headache with a probability of 0.5 but unlike medicine S, never occurred in combination with any other medicine, so that contrasting S with s’ identifies differences due to compound presentation. From a modeling perspective, an algorithm that relies solely on outcome variance to track uncertainty would only respond to medicines S and s’. In contrast, a computation that treats *both* confounding and stochasticity as sources of uncertainty should generate an increased signal to medicine C as well as S and s’: These divergent predictions are respectively instantiated by the error-driven and Bayesian causal model specified in the subsequent section.

On each trial of feedback-based learning (see Figure 1A), participants were presented with an individual allergy patient and were told either that the patient had not received any medicine or that a particular medicine, or combination of medicines, had been administered. Medicines were color coded and labeled accordingly (i.e., Medicine “B” was blue) with color assignments counterbalanced across participants. On the first screen, the allergy patient’s state was obscured by a question mark and participants were asked to indicate whether or not the patient had a headache, pressing the ‘‘Y’’ key for ‘‘yes’’ and the ‘‘N’’ key for ‘‘no”. After a prediction was made, there was a brief (2000ms) pause during which the word “wait” was displayed on the screen, followed by a screen revealing the allergy patient’s state (i.e., with or without headache).

**Figure 1.**
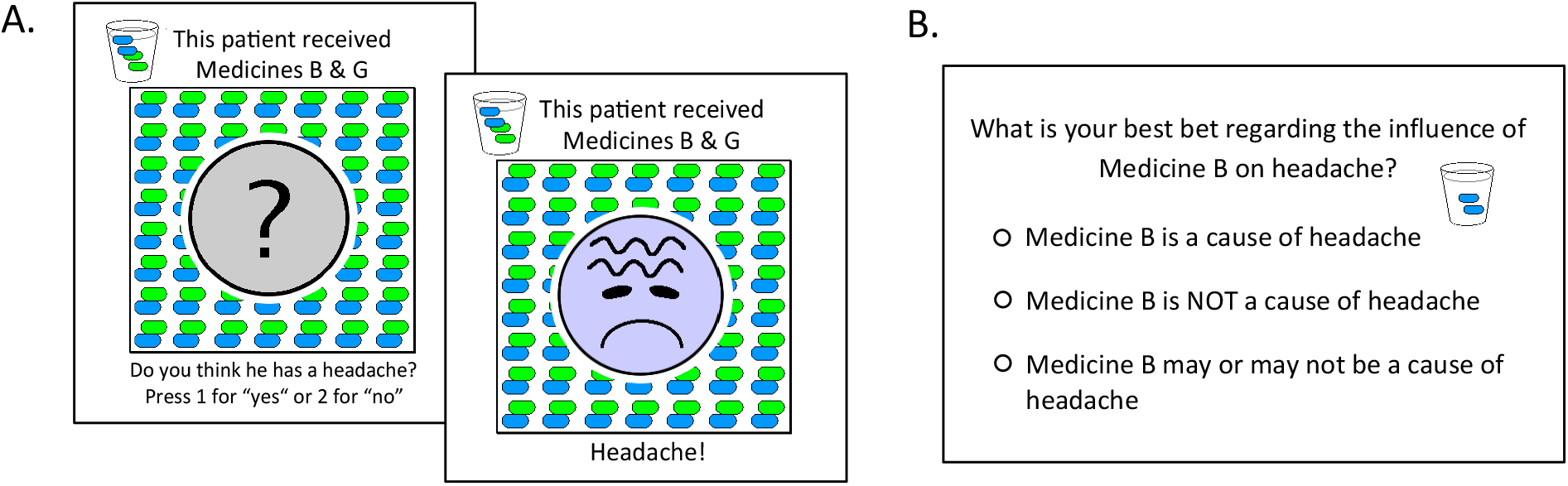
Task illustration. A. Prediction and outcome screens from a feedback-based learning trial. Screens were separated by a 2 sec. pause during which the word “Wait” (not shown in figure) was displayed. B. Causal query regarding the individual influence of a particular medicine on headache.

Each of 6 distinct trial types was presented 8 times in each of 4 sessions, for a total of 192 trials, with sessions separated by 2-minute breaks, during which the scanner was turned off. Each session was further divided into 4, non-delineated, blocks, with each type of trial occurring once in random order in each block. All trials were separated by a jittered 4-second ITI. In every other block, each trial (except for n-trials) was followed by a query regarding the individual causal influence of the target medicine present on that trial (i.e., C, D or S), or of medicine e or s’ on non-target trials. Specifically, participants were asked to choose between the following three options regarding the relevant medicine: “is a cause of headache”, “is not a cause headache” and an ambivalent “may or may not be a cause of headache” (Figure 1B). Categorical response options, rather than the rating scales commonly employed in the causal learning literature (Lu et al., 2008; Liljeholm & Cheng, 2009; Liljeholm, 2015) were used for two main reasons: First, because the limited number of response buttons in the scanner would have necessitated a scale slider, the initial positioning of which might potentially bias ratings, and second, to reduce movement due to back-and-forth scale scrolling.

### Computational models

Two formal accounts of predictive strength and uncertainty were implemented. First, an error-driven, associative learning algorithm updates predictive strength, on each trial, using the difference between the observed state of the outcome and the expected state based on all present cues:

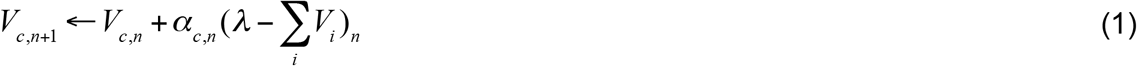

where *V* is the predictive strength of a particular cue, *c, ΣV_i_* is the summed strength of all cues present on trial *n*, and *λ* is the observed state of the outcome on that trial. The associability of cue *c* on trial *n, α_c,n_*, is defined as:

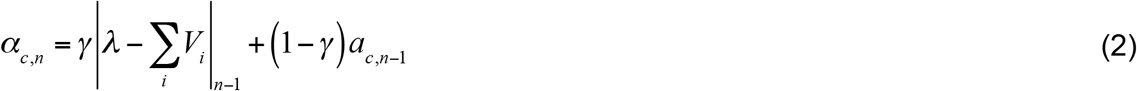

where *n-1* references to the previous trial involving cue *c*, and *γ* is a free parameter accounting for the weighting of that previous trial relative to preceding ones. In addition to scaling the influence of the prediction error on predictive strength, *α* is taken to reflect current levels of uncertainty about the cue’s influence (e.g., Esberg & Haselgrove, 2011). The strengths and uncertainties associated with each cue were initialized to 0.5. On feedback-based trials with target causes, which always occurred in compound with an alternative cause, the α of the target cause (always greater than or equal to that of its compound counterpart) was used to indicate uncertainty, while the summed strength, *ΣV*, was used to predict the outcome, and to compute the prediction error.

A second formal account is provided by a Bayesian Causal Model, in which reasoners make inferences over causal structures potentially responsible for the observed data (Griffiths & Tenenbaum, 2005; Lu et al., 2008; Liljeholm, 2015). Here, as illustrated in Figure 2, three possible graphs (*Graphs 0-2*) were defined for a candidate cause, *C*, an alternative cause, *A*, and a constantly present background cause, *B*, where a causal link to the effect exists for neither *C* nor *A* (*G_0_*), *C* only (*G_1_*), or both *C* and *A* (*G_2_*).

**Figure 2.**
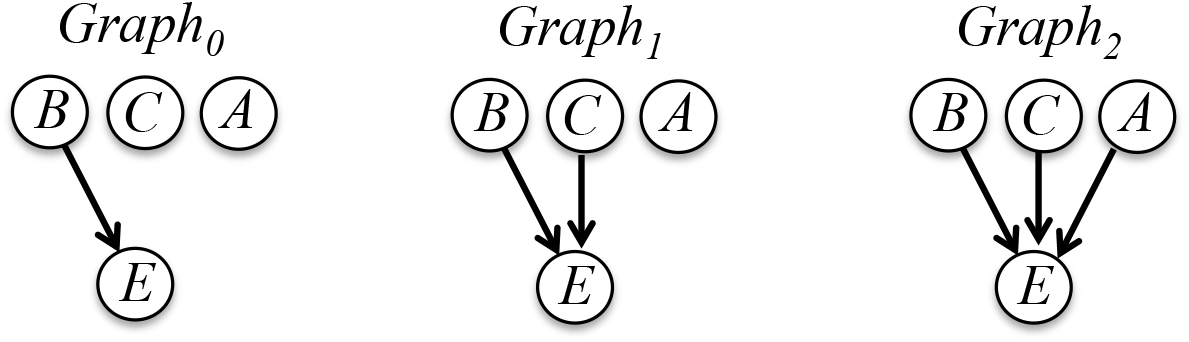
Possible causal structures involving a background cause, *B*, a target cause, *C*, an alternative cause, *A*, and an effect, *E*. A link to the effect always exists for *B*, and may exist for neither *C* nor *A* (*Graph_0_*), *C* only (*Graph_1_*), or both *C* and *A* (*Graph_2_*).

Each graph has a set of parameters *θ*, which are strengths *w_i_*, associated with causal links in each graph (i.e., *w_B_, w_C_* and *w_A_* for links associated with *B, C* and *A* respectively in *G_2_*). Sequential estimates of these parameters were modeled, for each candidate cause, using the smallest possible *focal set* (Cheng & Novick, 1992) – a set of events across which alternative causes can be assumed to occur with the same probability in the presence and absence of the relevant cause. Thus, estimates of the strength of the “no medicine” background cause was modeled using only n-trials in *G_0_*. Estimates of elemental (non-target) candidate causes s’ and e were respectively modeled based on trials with those causes and the “no medicine” trials (i.e., n- and s’± for candidate s’, and n- and e-for candidate e), using *G_1_*. Finally, for each target candidate cause, which always occurred in compound with some alternative cause, estimation was modeled given *G_2_*, using the trials relevant for the particular target cause (i.e., n- and Cc’ for target C; n-, e- and De+ for target D, and n-, e- and Se± for target S).

The likelihoods *P*(*d*|*θ, G_i_*) were computed using a noisy-OR parameterization (Cheng, 1997). Specifically, summarizing data *d* by contingencies *N*(*e,c,a*), the frequencies of each combination of the presence versus absence of the effect, target cause and alternative cause, the likelihood term for *G_2_* is:

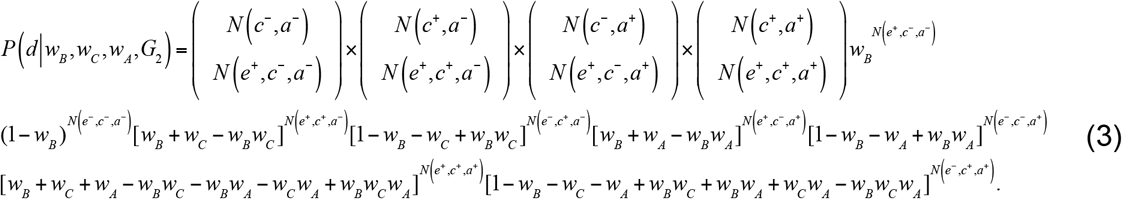

where 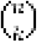 denotes the number of ways of picking *k* unordered outcomes from *n* possibilities. *N(c^+^)* indicates the frequency of events in which the target cause is present, with analogous definitions for the other *N(.)* terms. The likelihood terms for *G_0_* and *G_1_* are similarly specified, where frequencies *N(.)* are summed across the presence and absence of relevant events, and *w_i_=0* for any cause that does not have link to the effect in the relevant graph.

The marginal posterior distribution over strengths for a particular cause is obtained by applying Bayes’ rule, and integrating out the parameters for other causes in the graph, such that, for *G_2_*,

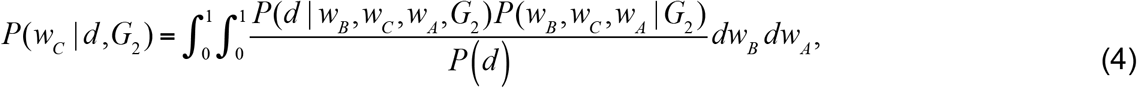

where *P(d|w_B_, w_C_, w_A_, G_2_)* is the likelihood term, *P(w_B_,w_C_,w_A_|G_2_)* refers to the prior probabilities of causal strength parameters, and *P(d)* is the normalizing term, denoting the probability of the observed data.

Uncertainty about the strength of a particular cause, the focus of the study, was modeled as the Shannon entropy *H(w_c_)* of its marginal posterior distribution *P(w_C_|d)*:

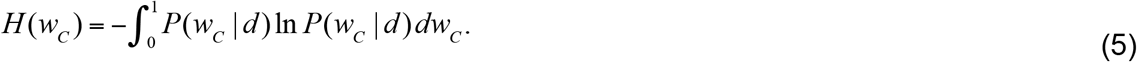

Point estimates of causal strength for each elemental and target cause were modeled as the mean of the relevant marginal posterior distribution (Lu et al., 2008; Liljeholm, 2015), given by:

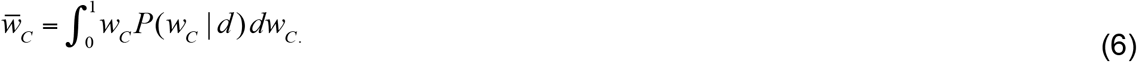

The model was sequentially implemented on a trial-by-trial basis, such that, for each cause, likelihoods were computed on each trial of feedback-based learning that yielded information relevant to that cause, using only the data point provided on that trial, with the posterior on the previous trial being used as the prior to obtain the posterior on the current trial. On the first trial in which data relevant to a particular candidate cause was presented, the priors were assigned independent uniform distributions. Finally, as an analog to the model-free prediction error, the Kullback-Leibler (KL) divergence between the prior (*Q*) and posterior (*P*) on each trial, commonly referred to as “Bayesian surprise” (Itti & Baldi, 2006) was computed as:

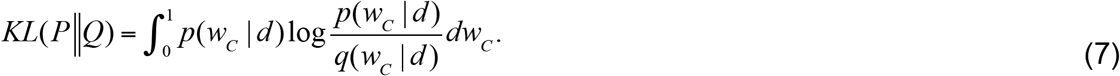

As noted, target causes only occurred in compound with alternative causes during feedback-based learning. On such trials, the entropy and KL divergence of the marginal distribution for the target cause was used to indicate uncertainty and surprise respectively, while predictions regarding the occurrence of the effect were generated using a noisy-OR integration of causal strengths.

Both models assumed that, during feedback-based learning, participants selected “Yes”/”No” responses to questions about whether headache would occur using probabilities generated by a softmax distribution, in which a free noise parameter controls the influence of predictive strength (i.e., the strength with which the medicine(s) present on a given trail predicted headache; noisy-logical *P(e|w_i_)* and *ΣV* in causal and error-driven models respectively) on choice behavior. An additional noise parameter controlled the influence of uncertainty (i.e., *α* and *H(w_c_)* in error-driven and causal models respectively) on the proportion of ambivalent “May or may not be a cause” judgments during queries about the individual influence of each medicine. The first two instances of each trial type, and the first causal judgment for each medicine, were excluded from model fitting to eliminate transient noise due to task adjustment, and to minimize the influence of assumptions about priors. Free parameters were fit to behavioral data by minimizing the negative log likelihood of observed responses for each individual and the Bayesian Information Criterion (BIC) was used for model selection.

The model-derived probabilities of uncertain “May or may not be a cause” judgments are plotted in Figure 3 for the causal and error-driven account of uncertainty respectively. Note that, as was foreshadowed in the description of the study design, the Bayesian causal model predicts high levels of uncertainty for both confounded, C, and stochastic, S and s’, causes, while the error-driven algorithm predicts that only stochasticity will generate high levels of uncertainty.

**Figure 3.**
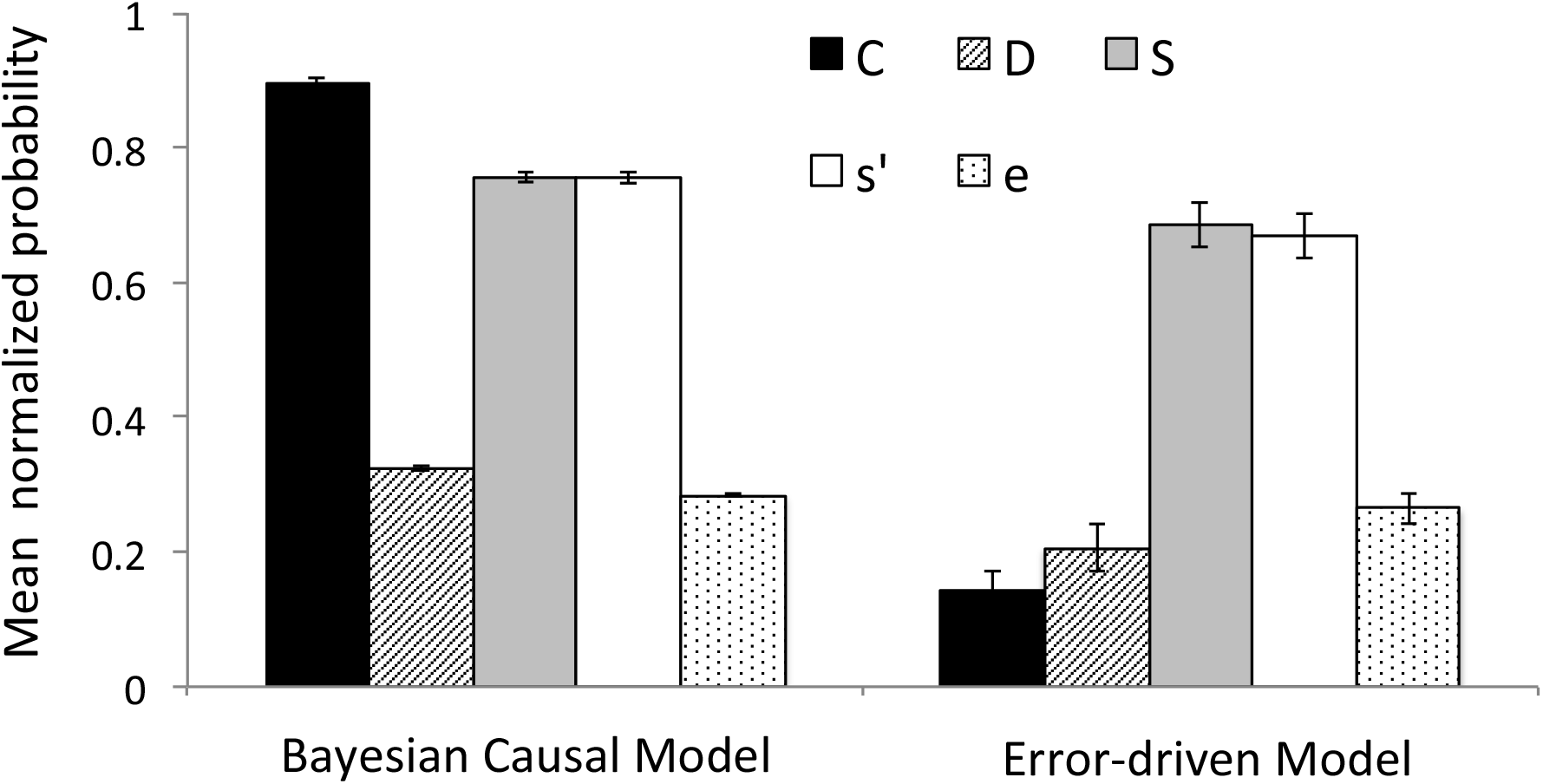
Mean (normalized) probabilities of uncertain “May or may not be a cause” judgments derived from the Bayesian causal model and error-driven algorithm respectively. Confounded, deterministic and stochastic target causes (C, D & S) occurred only in compound with alternative causes during feedback-based learning, while non-target causes occurred only individually (s’), or individually as well as in compound (e). Error bars=SEM.

### Imaging procedure and analysis

A 3 Tesla scanner (Phillips Achieva) was used to acquire structural T1-weighted images and T2*-weighted echoplanar images (repetition time = 2.65 s; echo time = 30 ms; flip angle = 90°; 45 transverse slices; matrix = 64 × 64; field of view = 192 mm; thickness = 3 mm; slice gap = 0 mm) with BOLD contrast. The first four volumes of images were discarded to avoid T1 equilibrium effects. All remaining volumes were corrected for differences in the time of slice acquisition, realigned to the first volume, spatially normalized to the Montreal Neurological Institute (MNI) echoplanar imaging template, and spatially smoothed with a Gaussian kernel (8 mm, full width at half-maximum). High-pass filtering with a cutoff of 128 s was used. All effects were reported at a whole brain FWE cluster corrected threshold of *p* < *0.05*, calculated using the Statistical nonParametric Mapping toolbox (SnPM13; http://warwick.ac.uk/snpm; Nichols & Holmes, 2002), with 5000 permutations, no variance smoothing and an uncorrected height threshold of *p* < *0.001*.

To address collinearity due to the shared prediction by causal and error-driven accounts that uncertainty increases with increased stochasticity, the two algorithms were analyzed using separate 1^st^ level General Linear Models (GLM), and were directly contrasted at the group level using Bayesian Model Selection. Specifically, for each participant, two GLMs were specified, one for the causal and one for the error-driven computational model. In each such GLM, three regressors respectively modeled 1) the onset of the prediction screen on each trial during feedback-based learning, 2) the onset of the outcome screen on each trial during feedback-based learning, and 3) the onsets of the screens soliciting judgments about the influences of individual medicines. In one GLM, prediction-screen onsets and causal judgment onsets were modulated by trial-by-trial error-driven estimates of uncertainty and strength, while outcome-screen onsets were modulated by trial-by-trial absolute values of the prediction errors. In the second GLM, prediction-screen and causal judgment onsets were modulated by trial-by-trial estimates of uncertainty and strength by the causal model while outcome-screen onsets were modulated by trial-by-trial estimates of Bayesian surprise. In both models, the first two instances of each trial type, and corollary causal judgments, were modeled by separate regressors, to eliminate transient noise due to task adjustment and reduce the influence of, non-modeled, prior beliefs. Moreover, in each model, two onset regressors respectively modeled the responses on feedback-based and causal query trials, three regressors indicated separate sessions, and six additional regressors accounted for the residual effects of head motion. Except for those indicating sessions and head motion, all regressors were convolved with a canonical hemodynamic response function. No orthogonalization was applied.

To directly contrast error-driven and causal accounts, BMS analyses was performed using a set of GLMs with the same onsets as those specified above, but with a single parametric modulator (e.g., error-driven uncertainty at prediction screen onsets) entered for each GLM. Thus, model comparisons isolated the relative contribution of a particular computational variable to neural activity. First-level Bayesian estimation procedure was used to compute a log-model evidence map for every subject and each GLM and inferences at the group level were modeled by applying a random effects approach (Rosa et al., 2010) at every voxel of the log evidence data. Group-level exceedance probability maps, reflecting the probability that one model is more likely than the other, were thresholded to identify voxels in which the exceedance probability was greater than 0.95. Classical inferences were then assessed using BMS masks, within which a particular computational variable was more likely than its competing counterpart, at an exceedance probability greater than 0.95: For example, significant effects of error-driven uncertainty during the prediction period of feedback-based learning trials were reported only for those regions in which error-driven uncertainty during this event period was more likely than causal uncertainty during the same period, according to the BMS.

### A pilot study on error-driven uncertainty

Previous research has implicated the anterior insula and dorsal medial frontal cortex in outcome stochasticity, using a range of decision-making tasks (Mohr et al., 2010; Volz et al., 2003; Huettel et al., 2005; Grinband et al., 2006). To ensure that such effects, used here as a benchmark against which to assess neural representations of uncertainty due to confounding, also emerge in our causal learning task, a pilot study (n=10) was conducted that was highly similar to that reported here, with the following exceptions: First, only stochasticity, not confounding, was manipulated across putative causes, second, both generative and preventive causal influences were included and, finally, judgments of individual causal influences were solicited at the end of, rather than throughout, each scanning session, and measured on a scale raging from −100 (strongly removes headache) to +100 (strongly produces headache). The error-driven model of uncertainty was implemented as described above, fit to behavior during feedback-based learning, and regressed against the BOLD data. A single region of interest (ROI) was constructed from the anterior insula and dorsal medial frontal cortex, using the “Willard” functional parcellation atlas (Richiardi et al., 2015). A power analysis performed with NeuroPower (http://neuropowertools.org; Durnez et al., 2015), using a screening threshold of z=2.3, revealed that a sample size of 16 would yield a power of 0.9 to detect effects of error-driven predictive uncertainty in this ROI at a Gaussian random field corrected alpha level of 0.05.

## Results

### Behavioral results

All statistical tests of behavioral data were planned comparisons, employing two-tailed t-tests, and were calculated using *n*=19. Confidence intervals (*CI*) and effect sizes (Cohen’s *d*) are reported for all comparisons. The distributions of causal judgments across response options are shown, with associated response times, for each putative cause in Table 1. Note that the proportion of ambivalent “May or may not be a cause” judgments (3^rd^ row in Table 1) were greatest for the confounded cause (C), intermediate, although still substantial, for the two stochastic causes (S and s’), and virtually absent for deterministic causes (D and e), exactly as predicted by the causal, but not the error-driven, model (cfr Figure 3; see Liljeholm, 2015 for similar results). Note also that the distribution of “Is a cause” and “Is not a cause” judgments differed markedly across stochastic and confounded causes, such that participants almost never select the latter option for the confounded cause, consistent with the normative increase in the likelihood that target cause C is causal given Cc+ trials, but distributed their responses fairly evenly across these two options for stochastic causes, reflecting, perhaps, the outcome on trials immediately preceding each judgment.

**Table 1.** Mean proportions of “Is a cause”, “Is not a cause” and “May or may not be a cause” judgments, and associated response times, with standard deviations, for target causes, C, D and S, and non-target causes, s’ and e’.

|  | C | D | S | s’ | e |
| --- | --- | --- | --- | --- | --- |
| *Proportions* |  |  |  |  |  |
| Is a cause | 0.36±0.33 | 0.84±0.25 | 0.38±0.25 | 0.40±0.27 | 0.02±0.03 |
| Is not a cause | 0.03±0.04 | 0.06±0.09 | 0.23±0.11 | 0.30±0.17 | 0.96±0.05 |
| May be a cause | 0.61±0.35 | 0.10±0.21 | 0.39±0.31 | 0.30±0.28 | 0.02±0.04 |
| *Response Times* |  |  |  |  |  |
| Is a cause | 2.92±1.88 | 2.52±1.23 | 4.03±3.62 | 2.82±2.59 | 2.86±2.11 |
| Is not a cause | 4.26±2.69 | 3.94±3.36 | 4.19±3.64 | 2.72±1.23 | 2.44±1.15 |
| May be a cause | 3.79±2.68 | 3.76±1.46 | 2.99±1.94 | 2.59±1.32 | 4.08±4.78 |

Planned comparisons revealed that the mean proportion of ambivalent causal judgments were significantly lower for the deterministic (D) target cause than for both confounded (C), t(18)=6.78, *p*<0.001, *CI*=[0.35 0.67], *d*=1.79, and stochastic (S), t(18)=4.09, *p*<0.001, *CI*=[0.14 0.44], *d*= 1.11, target causes. The difference between the confounded and stochastic cause was also significant, with the mean proportion of ambivalent judgments being greater for C than for S t(18)=2.65, *p*<0.05, *CI*=[0.0.05 0.40], *d*=0.68. Recall that all target causes, including S, occurred only in compound with alternative causes during feedback-based learning. The proportion of ambivalent judgments for the two stochastic causes, one always occurring in isolation (s’) and the other always in compound (S), was marginally significant, *p=0.08*.

Judgment response times, averaged across response options for each cause, were significantly faster for judgments about the deterministic target cause D than stochastic target cause S, t(18)=2.20, *p*<0.05, *CI*=[0.03 1.02], *d*=0.35, but did not differ significantly between D and C, nor between C and S, *p>0.1*. Moreover, response times did differ significantly across the two stochastic causes, being significantly slower for target cause S, which never occurred in isolation during feedback-based learning, than for cause s’, t(18)=3.42, *p*<0.005, *CI*=[0.29 1.21], *d*=0.52.

With respect to model fitting, the mean best fitting values of the free parameters for the error driven model were 0.35±0.27 for the weighting parameter, 3.25±0.95 for the prediction noise parameter, and 1.08±0.88 for the judgment noise parameter. The mean best fitting values of the free parameters for the causal model were 4.13±1.44 for the prediction noise parameter and 1.03±1.05 for the judgment noise parameter. The mean BIC was significantly lower, indicating superior performance, for the causal model (220.00±44.15) than for the error-driven model (237.09±35.58), t(18)=2.51, *p*<0.05, *CI*=[2.81 31.37], *d*=0.43.

### Neuroimaging results

All results reported below survived inclusive masking with voxels identified by Bayesian model selection analyses. A complete list of the non-masked significant effects of each model is provided in Tables 2 and 3. Bar graphs show effect sizes extracted from 8mm spheres centered on peak MNI coordinates, using rfxplot (Glascher, 2009).

**Table 2:** Non-masked significant, whole-brain corrected, neural effects of the causal model of uncertainty, strength and surprise, during causal judgments and feedback-based learning, with peak MNI coordinates and cluster sizes at p<0.001. R/L = Right/Left, (-) = negative correlation.

| Contrast | Region | Peak MNI | Cluster Size |
| --- | --- | --- | --- |
| Judgment Uncertainty | DMPFC | -8, 28, 46 | 1900 |
|  | R. IPG | 50, -52, 40 | 292 |
|  | Dorsolateral PFC | 42, 34, 30 | 201 |
| Prediction Uncertainty | Fusiform | 28, -72, -12 | 7195 |
|  | DMPFC | -10, 16, 52 | 1387 |
|  | R. Insula | -30, 18, 4 | 726 |
|  | L. Insula | 34, 30, -4 | 470 |
|  | Thalamus | 6, -22, 2 | 867 |
| Prediction Strength | Calcarine/Occipital | 18, -96, 4 | 236 |
| Prediction Strength (-) | Lingual Gyrus | 14, -62, -2 | 2232 |
|  | Superior Frontal Gyrus | -26, -8, 58 | 458 |
|  | Superior Parietal Lobule | -28, -50, 62 | 304 |
|  | Inferior Parietal Gyrus | -46, -26, 36 | 336 |
|  | Supp. Motor Area | 8, 2, 52 | 297 |
| Surprise | Right Insula | 36, 18, -2 | 652 |
|  | Left Insula | -28, 22, -6 | 482 |
|  | Dorsomedial PFC | 4, 46, 34 | 1146 |
|  | Left Dorsolateral PFC | -46, 4, 42 | 192 |
|  | Right Dorsolateral PFC | 44, 24, 30 | 325 |
| Surprise (-) | Supramarginal Gyrus | -50, -26, 38 | 1858 |
|  | Caudate | 18, 32, 0 | 505 |
|  | Mid Cingulum | -10, -30, 42 | 472 |
|  | Postcentral Gyrus | 50, -20, 48 | 2399 |
|  | Parahippocampal | 34, -32, -12 | 227 |
|  | Precuneus | 16, -52, 14 | 221 |

**Table 3:** Non-masked significant, whole-brain corrected, neural effects of the error-driven model of uncertainty, strength and surprise during feedback-based learning, with peak MNI coordinates and cluster sizes at p<0.001. R/L = Right/Left, (-) = negative correlation.

| Contrast | Region | Peak MNI | Cluster Size |
| --- | --- | --- | --- |
| Prediction Uncertainty | Calcarine/Lingual Gyrus | 14, -88, 0 | 3116 |
|  | Supp. Motor Area | 10, 20, 48 | 153 |
|  | Left Insula | -24, 22, 2 | 65 |
|  | Right Insula | 34, 28, -2 | 109 |
| Prediction Uncertainty (-) | Middle Temporal Gyrus | -46, -62, 20 | 341 |
|  | Middle Temporal Gyrus | -56, -22, -10 | 84 |
|  | Ventromedial PFC | -2, 30, -14 | 157 |
|  | Middle Temporal Gyrus | 46, -58, 22 | 64 |
| Prediction Strength | Lingual Gyrus | -20, -78, -10 | 2801 |
| Prediction Strength (-) | Superior Parietal Lobule | -18, -52, 64 | 407 |
|  | Supramarginal Gyrus | -60, -28, 30 | 390 |
|  | Precentral Gyrus | 26, -12, 50 | 240 |
|  | Supramarginal Gyrus | 58, -38, 28 | 728 |
|  | Middle Temporal Gyrus | -54, -60, 2 | 603 |
|  | Precentral Gyrus | -22, -14, 64 | 383 |
|  | Thalamus | -12, -10, -4 | 214 |
| Prediction Error | Right Insula | 38, 18, -4 | 912 |
|  | Left Insula | -40, 20, 4 | 945 |
|  | Precentral Gyrus | -42, -2, 40 | 713 |
|  | Anterior Cingulate | 4, 38, 30 | 2021 |
|  | Inferior Parietal Gyrus | -28, -52, 42 | 246 |
|  | Dorsolateral PFC | 44, 28, 26 | 885 |
| Prediction Error (-) | Middle Occipital Gyrus | -42, -74, 26 | 735 |
|  | Fusiform Gyrus | -34, -42, -12 | 253 |
|  | Postcentral Gyrus | 54, -26, 52 | 906 |

#### Uncertainty signals during causal judgments

During causal queries, as participants made judgments about the causal influences of individual medicines, selective and significant effects of the causal model of uncertainty emerged in the medial and lateral superior frontal gyrus and the right dorsolateral prefrontal cortex (Figure 4). Note that the effect size of BOLD responses (bar graphs in Figure 4) across putative causes were greatest for the confounded cause (C), intermediate for stochastic causes (S and s’), and smallest for deterministic causes (D and e), exactly as predicted by the causal, but not the error-driven, model (cfr Figure 3). To further probe these effects, response times (RTs) were included as a parametric modulator of judgment uncertainty in 1^st^ level, individual-subject, models, and as a between-subjects covariate in group-level analyses. Neither of these additional analyses revealed any modulatory influence by RTs on the neural effects of the causal model of judgment uncertainty. No effects of the error-driven measure of uncertainty during causal judgments survived whole-brain correction.

**Figure 4.**
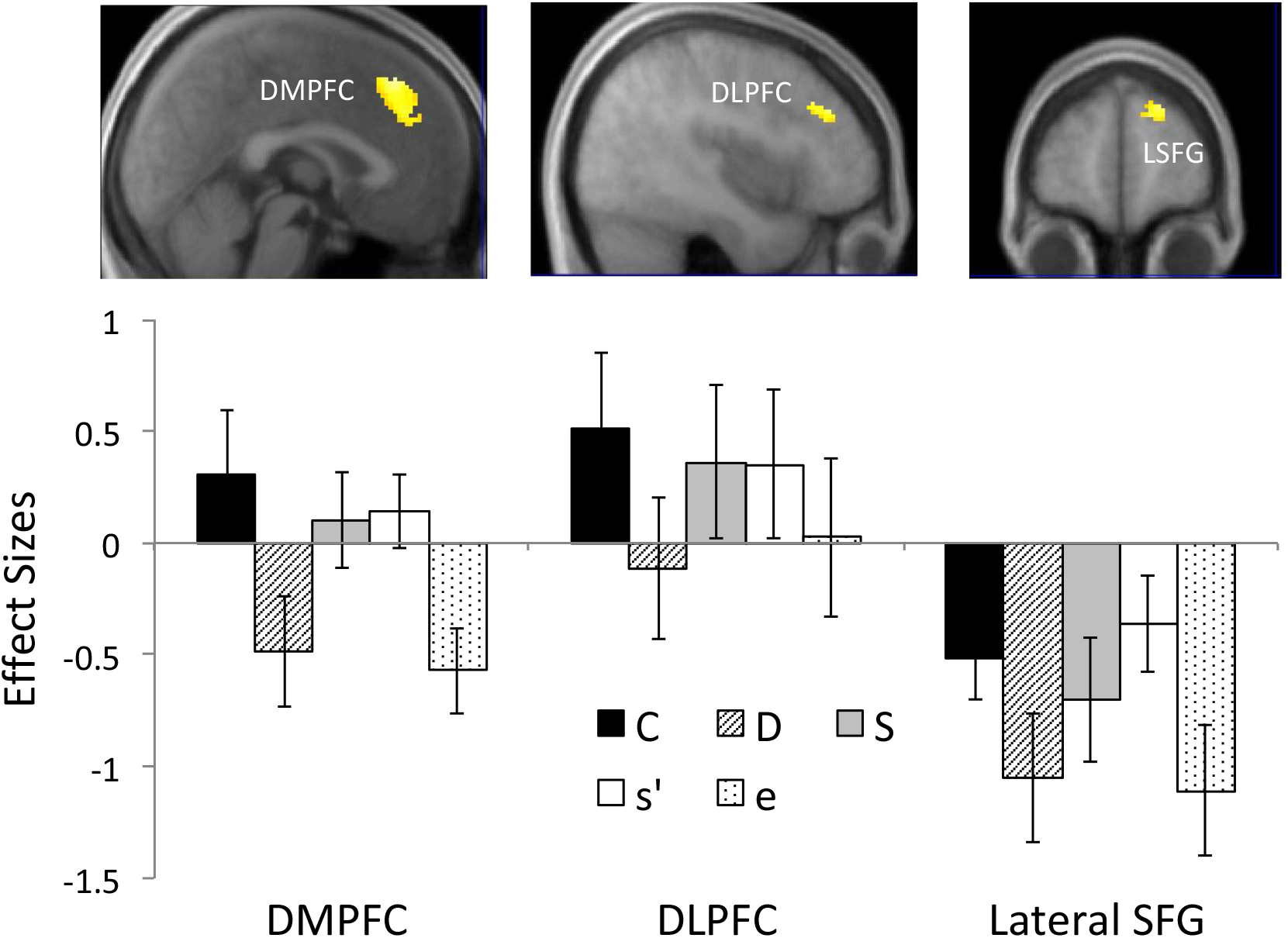
Selective effects of the causal model of uncertainty during judgments about the causal influences of individual medicines, showing effects in the dorsomedial and dorsolateral prefrontal cortex (DMPFC and DLPFC), and the lateral Superior Frontal Gyrus (LSFG). Bar plots show effect sizes extracted from spheres centered on peak coordinates for each queried medicine. Error bars=SEM.

#### Uncertainty signals during prediction

During the prediction period of feedback-based learning trials, the error-driven, but not causal, measure of uncertainty was significantly positively correlated with activity throughout the inferior occipital and lingual gyri, as well as with activity in the supplementary motor area and right anterior insula (Figure 5). Moreover, activity negatively correlated with error-driven predictive uncertainty was observed in the posterior middle temporal gyrus.

**Figure 5.**
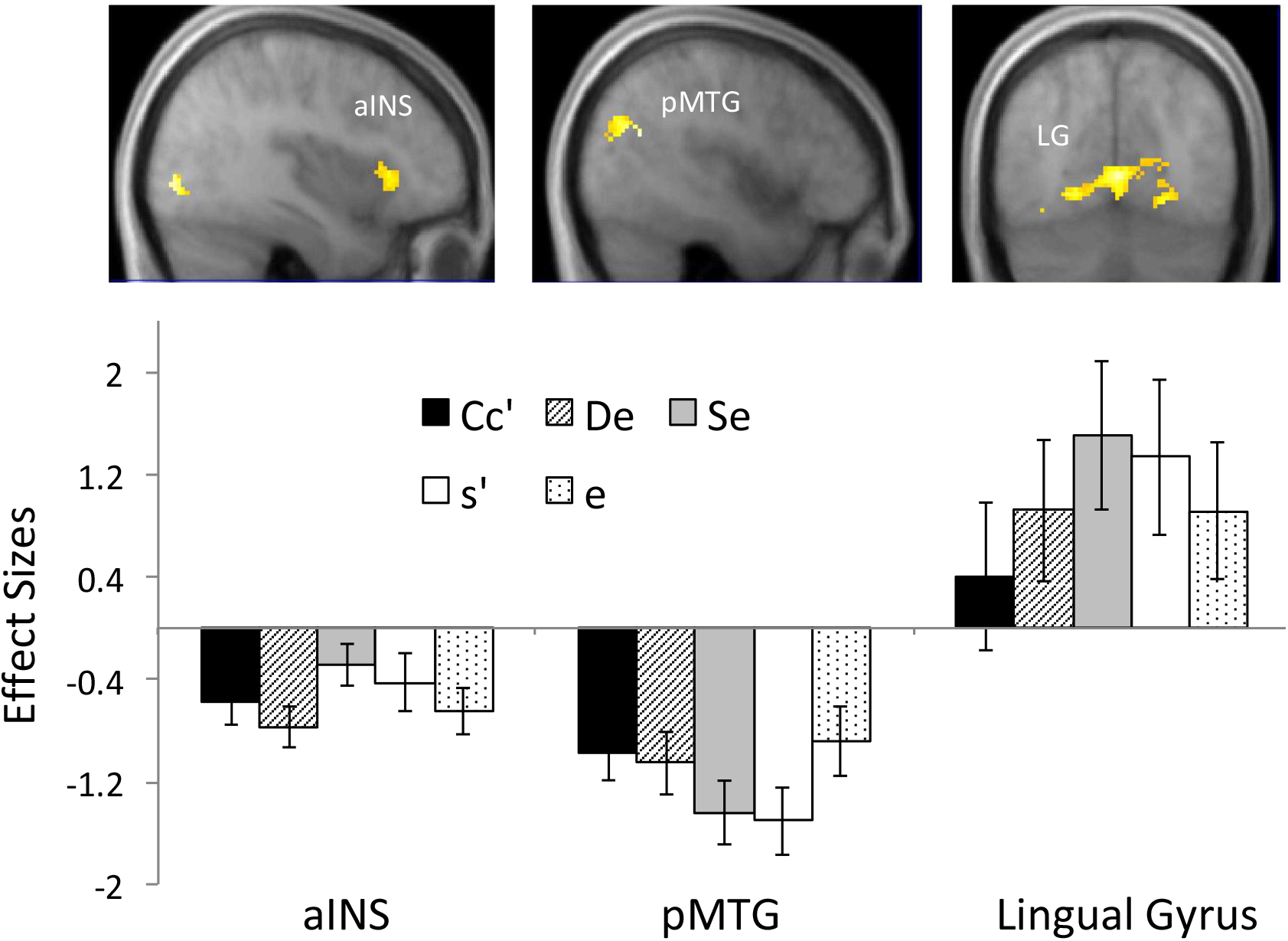
Selective effects of the error-driven model of uncertainty during the prediction period of feedback-based learning trials, showing effects in the right anterior insula (aINS), posterior middle temporal gyrus (pMTG) and lingual gyrus (LG). Bar plots show effect sizes extracted from spheres centered on peak coordinates for each type of medicine trial. Error bars=SEM.

These effects of the error-driven account of uncertainty appear relatively robust, as they closely mirror those from the pilot study, in which, at an uncorrected threshold of 0.005, extensive (cluster size > 100) effects of error-driven predictive uncertainty emerged in the dorsal medial frontal cortex, bilateral anterior insula, and lingual gyrus during feedback-based learning. In contrast to the widespread effects of error-driven predictive uncertainty during feedback-based learning, selective and significant effects of the causal measure of uncertainty emerged only in the right dorsal thalamus.

#### Strength & surprise signals

While uncertainty in causal inferences, due to confounding or stochasticity, is of primary interest here, estimates of the strength with which a cause, or combination of causes, predicts the effect, and of the surprise at the outcome on each trial of feedback-based learning, were also modeled for completeness. During the prediction period of feedback-based learning trials, the causal measure of predictive strength was selectively and significantly positively correlated with activity in the calcarine and superior occipital gyrus, as well as negatively correlated with activity in the left lateral superior frontal and precentral gyri (Figure 6). In contrast, a selective and significant negative correlation with the error-driven measure of predictive strength emerged in the posterior MTG. No significant effects of either the causal or error-driven account of predictive strength were found during causal judgments.

**Figure 6.**
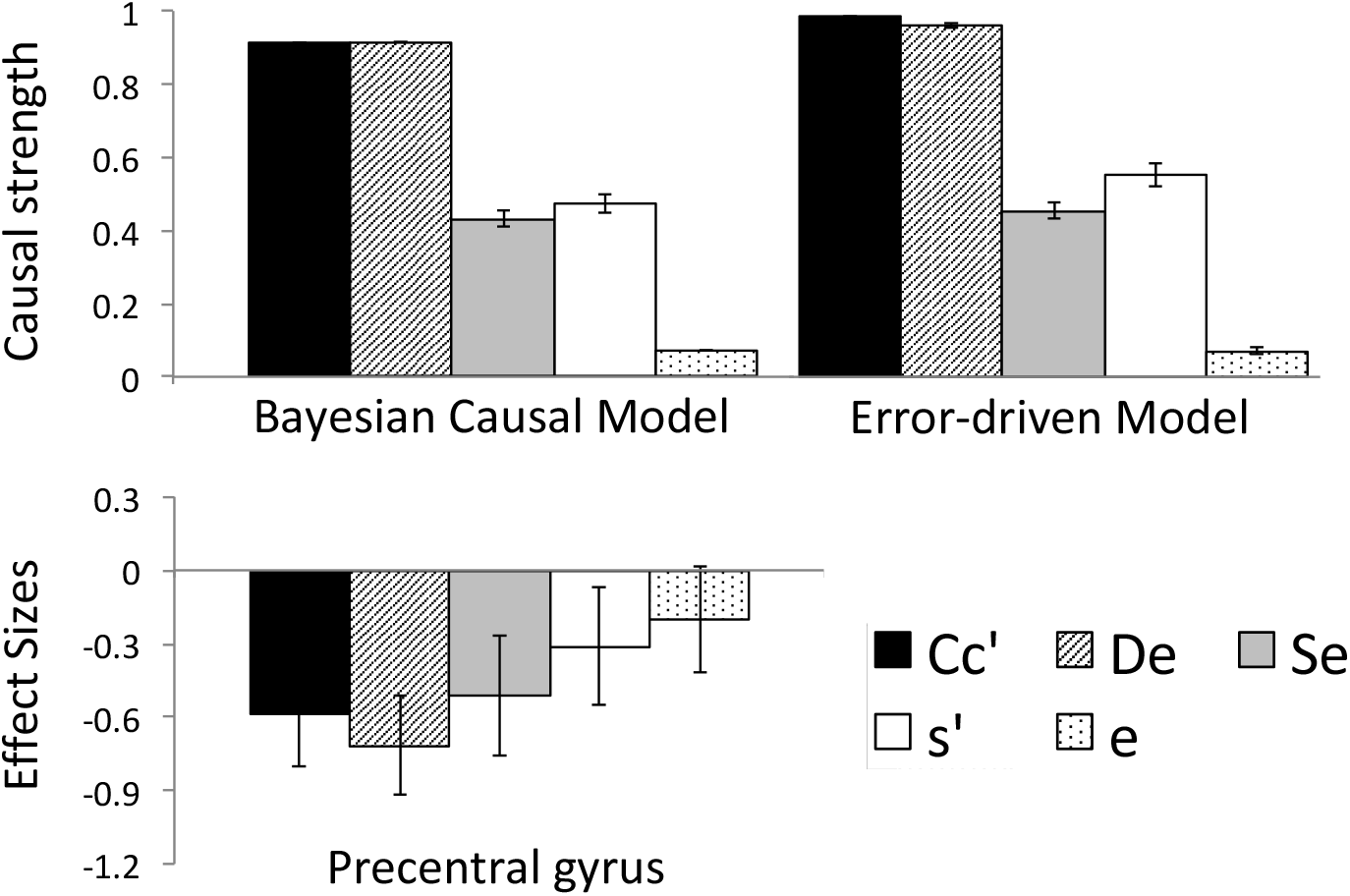
Strength predictions derived from the causal and error-driven model (top), together with effect sizes (bottom) extracted from spheres centered on peak coordinates in the precentral gyrus, for each type of medicine trial during feedback-based learning. Error bars=SEM.

For measures of surprise during the outcome period of feedback-based learning trials, the absolute prediction error generated by the error-driven account was selectively and significantly positively correlated with activity in the bilateral anterior insula, the medial and lateral dorsal prefrontal cortex and the inferior parietal gyrus. In addition, significant negative correlations with this measure were identified in the middle occipital gyrus, fusiform gyrus, the superior parietal lobule, precentral gyrus and calcarine sulcus. In contrast, a selective and significant negative correlation with the causal measure of (Bayesian) surprise was observed in the anterior caudate.

## Discussion

Sensitivity to the “no confounding” constraint – a requirement that putative causes do not co-vary, fundamental to philosophical and statistical perspectives on causation – has been reliably demonstrated in a range of behavioral judgments by children as well as adults (Spellman, 1996; Kushnir and Gopnik, 2005; Meder et al., 2006; Schulz and Bonawitz, 2007; Schulz et al., 2007; Liljeholm, 2015). The current study used fMRI and cognitive computational modeling to investigate neural substrates mediating uncertainty associated with causal confounding. While being scanned, participants studied and judged the influences of various putative causes with confounded or non-confounded, deterministic or stochastic, influences. During judgments requiring generalization of causal knowledge from a feedback-based learning context to a transfer probe, activity in the dorsomedial prefrontal cortex (DMPFC) was better accounted for by a Bayesian causal model, sensitive to both confounding and stochasticity, than a purely error-driven algorithm, sensitive only to stochasticity.

The DMPFC has been implicated in encoding uncertainty across a variety of tasks (Volz et al., 2003; Grinband et al., 2006; Xue et al., 2008; Michael et al., 2015). For example, scanning participants with fMRI as they classified stimuli according to a variable category boundary, Grinband et al. (2006) found that activity in the DMPFC scaled with the proximity of a stimulus to a category boundary. An important aspect of this and other demonstrations of the involvement of the DMPFC in uncertainty is that the level of uncertainty is directly related to the degree of experienced errors: that is, the greater the uncertainty on a given trial, the more likely it is that the prediction on that trial will be incorrect, as indicated by explicit feedback. In contrast, during feedback-based learning in the current task, the outcome on confounded trials was deterministic. Consequently, the increased activity in the DMPFC, as participants generated judgments about confounded cause, cannot be attributed to a history of errors. Instead, these results suggest that the DMPFC encodes a more abstract representation of uncertainty, divorced from the immediate consequences of choice, and irrespective of whether the specific source of uncertainty is confounding or stochasticity.

Several neuroeconomic studies (Hsu et al., 2005; Krain et al., 2006; Huettel et al., 2006; Bach et al., 2009; Levy et al., 2010; Pushkarskaya et al., 2015) have contrasted gambles involving stochasticity with “ambiguous” gambles, for which outcome probabilities are completely unknown, with some finding greater activity in the DMPFC in response to ambiguity than stochasticity (e.g., Hsu et al., 2005). One might argue that, in the current study, the influence of the confounded medicine is unknown, just as the outcome probabilities of ambiguous gambles in previous studies were unknown. But what does it mean to “know” something? Critically, all target medicines occurred only in compound with alternative medicines during feedback-based learning: consequently, when presented individually during causal queries, each was equally novel and, presumably, equally unknown. In other words, differences in ambiguity between confounded and other target causes cannot be attributed to a generalization decrement based on changes in stimulus features. Moreover, ambiguity due to small samples and uncertainty due to confounding call for very different plans of exploratory action: while in former case, repeated observation of the same stimulus can resolve the uncertainty, the latter case requires an intervention that unconfounds the stimulus configuration. Further work is needed to assess the overlap between neural representations of confounding and ambiguity.

In addition to stochasticity and ambiguity, the DMPFC has been heavily implicated in cognitive control (Ridderinkhof et al., 2004; Fellows & Farah, 2005; Egner, 2009; Taren et al., 2011; Shenhav et al., 2013), raising the question of whether the currently observed effects in this region might reflect similar processes. In particular, judgments about individual causal influences may be associated with efforts to retrieve instances of the queried cause occurring in isolation from episodic memory – a search that is futile, and thus presumably more effortful, for confounded causes. However, such retrieval-induced control demands are unlikely to account for the current effects in the DMPFC for several reasons: First, effort associated with episodic memory retrieval does not address responses to stochasticity – in contrast, the causal uncertainty account explains why DMPFC responses are greater for *both* confounded and stochastic causes, relative to non-confounded and deterministic causes. It may be argued, of course, that stochasticity elicits additional control processes mediated by the DMPFC, such as, for example, response conflict (Botnivick et al., 2001; Ridderinkhof et al., 2004; Kerns et al., 2004). Recall, however that *none* of the target causes (C, S and D) were presented in isolation during feedback-based learning and that, consequently, *all* target causes should involve some level of searching for non-presented information during causal judgments. If DMPFC activity reflected this search process as well as response conflict due to stochasticity, one would expect a difference between neural responses to causes S and s’, both of which are stochastic, but only one of which, s’, was presented in isolation during feedback-based learning, eliminating the need to search for an un-experienced memory. Notably, while RTs (Table 1), commonly used as a measure of cognitive load, are indeed consistent with the notion of an increased control demand for cause S over s’, the DMPFC responses (Figure 4) are not. Finally, and perhaps most conclusively, the inclusion of RTs as a parametric modulator of judgment uncertainty in 1^st^ level models, and as a covariate in group analyses, did not have any impact on the neural effects of model-based judgment uncertainty. Taken together, these aspects of the results make causal uncertainty the more parsimonious explanation of the DMPFC effects observed here.

As with the DMPFC and DLPFC, the anterior insula has been implicated in outcome stochasticity, or “risk”, in several previous neuroimaging studies (e.g., Volz et al., 2003; Huettel et al., 2005; d’Acremont et al., 2013; Preuschoff et al., 2008; Critchley et al., 2001). Here, the dorsal anterior right insula was selectively recruited by the error-driven model of uncertainty during the prediction period of feedback-based learning trials. Plots of effect sizes for each type of trial (in Figure 5) suggest that these responses were indeed driven by stochasticity–based uncertainty about the state of the outcome on any given trial. Moreover, the fact that, unlike the DMPFC, activity in the anterior insula did not scale with either an error-driven or causal measure of uncertainty during judgments about individual causal influences indicates that representations in this region pertain primarily to the immediate consequences of choice, rather than generalizable knowledge. Consistent with this interpretation, the bilateral anterior insula also responded selectively to stochastic conditions during the outcome period of feedback-based learning trials, suggesting that this region is recruited by both the anticipation and detection of errors. While, as noted, it is difficult to discern based on the current results whether the unsigned error signals observed in the anterior insula are Bayesian or model-free, they are consistent with a substantial literature implicating this region in expectancy violations across a wide range of tasks (Klein et al., 2007; Metereau et al., 2012; Allen et al., 2016; Bastin et al., 2016).

Selective effects of the error-driven account of predictive uncertainty were also identified throughout large portions of the posterior lingual gyrus. In a recent neuroimaging study, Causse et al., (2013) assessed decision making under uncertainty using an aviation task in which participants had to either land a plane or abort landing, given certain (100% or 0%) vs. uncertain (50%) information regarding landing conditions. Consistent with the current results, they found greater activity in the posterior lingual gyrus during high than during low uncertainty conditions. Our results are also consistent with those of Payzan-LeNestour et al. (2016), who found that the lingual gyrus scaled specifically with uncertainty due to outcome stochasticity, as opposed to uncertainty due to changes in the statistical structure of the environment or due to ambiguity. Payzan-LeNestour et al. argued that the lingual gyrus might not typically emerge in studies assessing outcome stochasticity because those studies did not dissociate stochasticity from other uncertainty components. A similar case can be made here, that the inclusion of an alternative, generative, model that captures additional aspects of uncertainty may have increased sensitivity to detect stochasticity-specific signals in the lingual gyrus.

An important aspect of the current study is the direct comparison of a Bayesian inference model with an error-driven, model-free, algorithm. While not uncommon (Courville et al, 2006; Hampton et al., 2006; Payzan-LeNestour & Bossaerts, 2011; Prevost et al., 2013), the contrasting of these quite different computational frameworks somewhat complicates interpretation. In particular, it raises the question of whether selective effects of the causal account of uncertainty indeed reflect sensitivity to confounding, or instead some extraneous, perhaps incidental, feature of the different modeling frameworks. Here, a significant clue is provided by neural and behavioral responses to each queried putative cause, shown in Figure 4 & Table 1 respectively, which clearly correspond to the increased uncertainty associated with confounded and stochastic causes predicted by the causal model (cfr Figure 3). It is worth noting that, far from incidental, the failure of the error-driven account to predict sensitivity to confounding is intrinsic to the model-free approach, due to its lack of a representation of the independence of causal influences. Critically, a configural solution (e.g., Pearce, 1987; 2002), such that individually queried causes are treated as non-overlapping with the compounds in which they occur during feedback-based learning, would fail to account for both behavioral and neural differences between confounded and deterministic target causes, both of which occurred only in compound with alternative causes during feedback-based learning.

While the effects of causal uncertainty in the DMPFC likely reflect sensitivity to causal confounding, other neuroimaging results are clearly unrelated and, in some cases, largely incidental to the specific model implementation. In particular, as can be seen in Figures 6 and 7, the ordinal mean values derived from causal and error-driven accounts across trial types during feedback-based learning are identical for both strength and surprise measures, suggesting that model-selection may reflect more granular differences. For example, because of the *Σ* term in Equations 1 and 2 of the error-driven account, the variance in the occurrence of the outcome across e-, De+ and Se± trials results in a constantly fluctuating strength, and persistent prediction errors, even on deterministic e- and De+, trials. In contrast, in the implementation of the causal model, such non-conditional variance is eliminated by the use of a focal set (e.g., Cheng & Novick, 1992; Spellman, 1996). Behaviorally, the focal set assumption is strongly supported by the lack of variability in judgments about non-target cause e. This cause, which is paired with the outcome on half of trials in which it occurs, but never when occurring in isolation, is almost exclusively judged to be non-causal, suggesting that participants conditioned their inferences about its influence on trials across which alternative causes could be assumed to be held constant. Nonetheless, at the neural level, several regions may instead have been tracking the unconditional outcome variance computed by the error-driven model.

**Figure 7.**
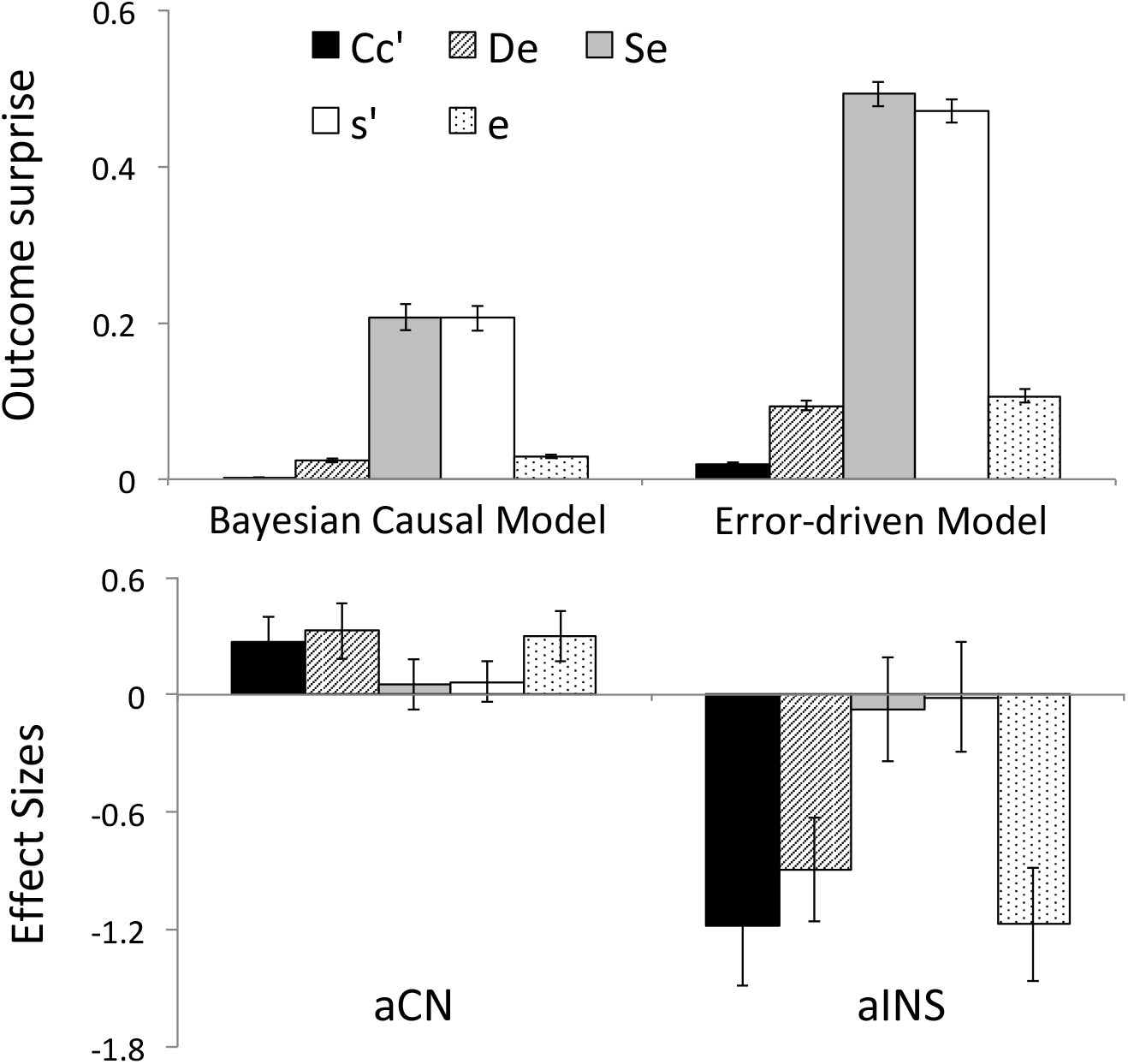
Outcome surprise signals derived from the causal and error-driven model (top), together with effect sizes (bottom) extracted from spheres centered on peak coordinates in the anterior caudate nucleus (aCN) and anterior insula (aINS), for each type of medicine trial during feedback-based learning. Error bars=SEM.

As the mantra “covariation does not equal causation” implies, requirements for causal induction extend far beyond mere statistical regularity. Countless studies have demonstrated neural and behavioral effects of error-driven uncertainty, whether due to stochasticity, insufficient samples or changes in the statistical structure of the environment. The current study reveals that, given violation of a basic boundary condition on causal inference, and in spite of the same constraints on perceptual generalization, differences in neural representations of uncertainty can emerge across equally deterministic, static and well-sampled contingencies. These results contribute to a growing literature on the role of generative models in neural representations of uncertainty, and shed light on a possible neural implementation of domain general, a priori, constraints on causal induction.

## Acknowledgements

This work was supported by a start-up fund from the University of California Irvine to M.L.

